# Akt is required for artery formation during embryonic vascular development

**DOI:** 10.1101/2020.06.04.134718

**Authors:** Wenping Zhou, Emma Ristori, Liqun He, Joey J Ghersi, Sameet Mehta, Rong Zhang, Christer Betsholtz, Stefania Nicoli, William C. Sessa

**Affiliations:** Department of Cell Biology, Yale University School of Medicine, New Haven, CT 06511, USA; Vascular Biology & Therapeutics Program, Yale University School of Medicine, New Haven, CT 06520, USA; Department of Pharmacology, Yale University School of Medicine, New Haven, CT 06510, USA; Yale Cardiovascular Research Center, Department of Internal Medicine, Section of Cardiology, Yale University School of Medicine, New Haven, CT 06511, USA; Department of Genetics, Yale University School of Medicine, New Haven, CT 06510, USA; Department of Immunology, Genetics and Pathology, Rudbeck Laboratory, Uppsala University, Dag Hammarskjölds väg 20, SE-751 85 Uppsala, Sweden; ICMC (Integrated Cardio Metabolic Centre), Karolinska Institutet, Novum, Blickagången 6, SE-141 57 Huddinge, Sweden; Department of Life Sciences, University of Siena, 53100 Siena, Italy

**Keywords:** zebrafish embryo, vascular development, endothelial cells, artery identity, akt signaling, FOXO1, Notch

## Abstract

One of the first events in the development of the cardiovascular system is morphogenesis of the main embryonic artery, the dorsal aorta (DA). The DA forms via a conserved genetic process mediated by the migration, specification, and organization of endothelial progenitor cells into a distinct arterial lineage and vessel type. Several angiogenic factors activate different signaling pathways to control DA formation, however the physiological relevance of distinct kinases in this complex process remains unclear. Here, we identify the role of Akt during early vascular development by generating mutant zebrafish lines that lack expression of *akt* isoforms. Live cell imaging coupled with single cell RNA sequencing of *akt* mutants reveal that Akt is required for proper development of the DA by sustaining arterial cell progenitor specification and segregation. Mechanistically, inhibition of active FOXO in *akt* mutants rescues impaired arterial development but not the expression of arterial markers, whereas Notch activation rescues arterial marker expression. Our work suggests that Akt activity is critical for early artery development, in part via FOXO and Notch-mediated regulation.

## INTRODUCTION

The cardiovascular system is essential to supply oxygen and nutrients to vital organs and tissues and is the first organ to function during vertebrate development. The dorsal aorta (DA) primordium is the main embryonic blood vessel that forms via endothelial progenitor cells (or angioblast) migration from the lateral posterior mesoderm, acquire arterial and venous gene expression programs, and then segregate to form the posterior cardinal vein (PCV) (Herbert et al., 2009a; Lawson and Weinstein, 2002a). With the formation of the vascular lumen and the initiation of the blood circulation, the DA and PCV acquire functional differences in vascular diameter, extracellular matrix and smooth muscle cell coverage (dela Paz and D’Amore, 2009; Lawson and Weinstein, 2002a) that are maintained into adulthood. Importantly, the molecular cues that dictate morphogenesis of the DA and PCV are conserved amongst vertebrates (Gore et al., 2012; Isogai et al., 2001).

The Ser and Thr kinase Akt, is a key element of the phosphotidylinositol 3-kinase (PI3K) /Akt signaling pathway and regulates the hallmarks of tissue formation such as growth and survival. Akt activation involves the phosphorylation of multiple substrates signaling network which enable Akt to exert diverse downstream effects from the activation of an individual upstream growth factor signaling. For example, during the formation of the DA, PI3K-Akt signaling is implicated in transducing multiple Vascular Endothelia Growth Factor a (Vegfa)-dependent cell behaviors, such as migration, differentiation and proliferation of endothelial cells. (Ackah et al., 2005; Liu et al., 2008; Phung et al., 2006; Zhu et al., 2013). Importantly, how Akt-substrate activation is involved in this multi phased process remain not fully understood.

Vertebrates encode three Akt isoforms: Akt1, Akt2, and Akt3 and these unique gene products are highly homologous between species. The Akt-signaling module is implicated in arteriogenesis by exerting an inhibitory influence on Raf-Erk-signaling (Ren et al., 2010) or in venogenesis via phosphorylation of the COUP TFII, a key driver of venous fate (Chu et al., 2016). However mice globally lacking Akt-1, the main isoform expressed in endothelial cells, are viable, smaller in size, with reduced angiogenesis in the placenta, retina and limbs (Ackah et al., 2005; Ha et al., 2019; Schleicher et al., 2009; Yang et al., 2003) whereas the conditional loss of Akt-1 in the endothelium demonstrates reduced post-natal arteriogenesis and sprouting of retinal (Lee et al., 2014) and coronary vessels (Kerr et al., 2016). Thus, the genetic loss of Akt1 or other Akt isoforms in vertebrate has not established clear roles for Akt during the early DA and/or PCV development.

To assess the role of Akt in the early embryonic vascular development, we generated a novel loss of function model in zebrafish. The external development and optical transparency of zebrafish embryos facilitate the temporal dissection of vascular defects that are difficult to assess in other vertebrates such as mice. Using CRISPR/Cas9, we mutated all four *akt* genes encoding all three Akt isoforms and crossed them to a reporter line to image arterial and venous structures during development. Live cell imaging coupled with single cell RNA sequencing analysis of endothelial cells reveal that Akt is required for arterial specification, and arteriovenous (A-V) segregation but not for progenitor migration, during the initial formation/remodeling of the DA. Mechanistically, our data suggest that the loss of Akt activates FOXO and reduces Notch dependent gene expression in arterial endothelium. Inhibition of FOXO1 in Akt deficient embryos increases DA development, whereas activation of Notch in these embryos enhances arterial specification. Thus, Akt controls two key transcription factors to coordinately govern DA morphology and arterial specification.

## RESULTS

### Embryonic loss of Akt results in artery-vein malformations

To investigate the role of Akt signaling in early embryonic vascular development, we generated a complete *akt* loss-of-function model in zebrafish. In zebrafish, Akt1 is encoded by *akt-1*, Akt2 is encoded by *akt-2*, and Akt3 is encoded by *akt-3a* and -3*b*. Of these four genes, *akt1* showed the highest expression in endothelial cells isolated from zebrafish (Figure S1A), consistent with observations in mice (Lee et al., 2014).

To establish a complete *akt* loss-of-function model, and avoid genetic compensation, we used CRISPR/Cas9 (Moreno-Mateos et al., 2015) to mutate all four akt genes (Figure 1A). We selected loss-of-function founders and established a quadruple homozygous mutant (akt^Δ/Δ^). Compared to wild-type (WT), akt^Δ/Δ^ embryos had an ∼80% reduction in the mRNA levels of each *akt* isoform (Figure 1B). akt^Δ/Δ^ embryos were not viable to adulthood, thus we established an akt^1Δ/Δ, 3aΔ/Δ, 3bΔ/Δ, 2 Δ/+^ which we named “mixed akt^Δ/Δ^”. The levels of total and phosphorylated Akt were significantly reduced in mixed akt^Δ/Δ^ relative to WT embryos (Figure 1B).

**Figure. 1.**
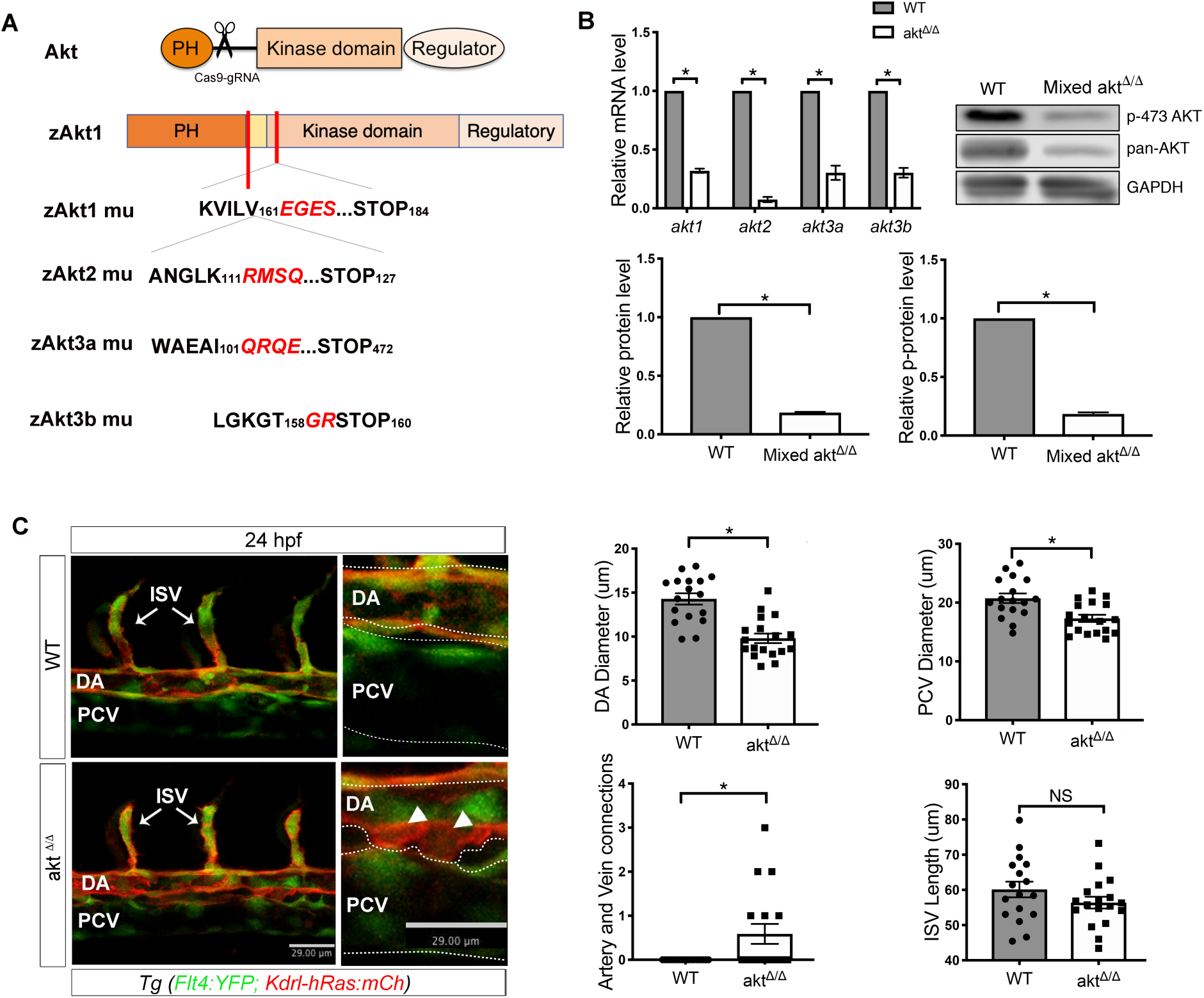
*akt*^*Δ/Δ*^ embryos exhibit decreased DA diameter and abnormal arterial and venous connections. (A) Schematic of the domain structure of Akt isoforms and sequence alignment of wild-type (zAKt1) and mutant (mu) zebrafish lines. (B) qRT-PCR showing WT and mutant mRNA levels in WT and quadruple mutant *akt*^*Δ/Δ*^ embryos on a *Tg (flt4:YFP; kdrl:hRas-mCherry)*^*hu4881;s896*^ at 4 days post fertilization (dpf). Expression levels were normalized to the WT from three biological replicates. Western blot analysis of WT and mixed *akt*^*Δ/Δ*^ embryos in *Tg (flt4:YFP; kdrl:hRas-mCherry)*^*hu4881;s896*^ from three biological replicates. *p< 0.05. (C) Confocal lateral view live images of WT and *akt*^*Δ/Δ*^ embryos at 24 hpf trunk in *Tg (flt4:YFP; kdrl:hRas-mCherry)*^*hu4881;s896*^. White arrowheads indicate the abnormal artery and vein connection in the representative images. Bar plots on right show in WT, and *akt*^*Δ/Δ*^ quantification. The embryos were placed in the lateral view with dorsal side up and anterior to left. Data are mean ± SEM for DA and PCV diameters and artery and vein connections; *p< 0.05. DA=Dorsal Aorta, PCV=Posterior Cardinal Vein, ISV=intersegmental vessels.

We assessed the overall morphology of akt^Δ/Δ^ embryos at early stages of development. Compared to WT, akt^Δ/Δ^ embryos exhibited a reduction in pigmented cells at 48 hours post fertilization (hpf) (Figure S1B) consistent with the chemical inhibition of Akt signaling (Ciarlo et al., 2017). Moreover, akt^Δ/Δ^ had normal body and head size (Figure S1B) and were viable until 6 days post fertilization (dpf), enabling the analysis of specific vascular patterns in during early development.

We crossed mixed akt^Δ/Δ^ line to a transgenic reporter line *flt4:YFP; kdrl:hRas-mCherry*^*hu4881;s896*^ to enable the analysis of arteries (mCherry+) and veins (YFP+/mCherry+) during embryonic development (Figure 1C). We performed live imaging of the vasculature at 24 hpf and found that akt^Δ/Δ^ embryos had decreased diameters of the endothelium of dorsal aorta (DA) and posterior cardinal vein (PCV) compared to WT embryos (Figure 1C images on left and quantified on right). The lengths of the Intersegmental vessels (ISV) that sprout from the DA appeared similar in both strains at this developmental time point. However, *akt*^*Δ/Δ*^ embryos showed ectopic connections between the DA and PCV (Figure 1C, arrowheads in magnified image and quantified on right) suggesting a lack of proper artery-vein segregation. These data suggest that Akt signaling input is necessary for proper patterning of the main embryonic blood vessels but dispensable for vascular cell migration of ISVs at 24 hpf.

### Loss of Akt impairs arterial specification

To investigate the molecular mechanisms that underline the vascular defects in embryos lacking Akt, we used single cell RNA sequencing analysis (scRNA-seq) of FAC-sorted endothelial cells (*kdrl*+ mCherry) from WT and mixed akt^Δ/Δ^ reporter embryos at 24 hpf. Mixed akt^Δ/Δ^ embryos showed substantially loss of Akt activity and recapitulated the same vascular defects of akt^Δ/Δ^ full mutants (Figure 1B and S1C).

Artery and vein formation during development generates cellular diversity in endothelial cells, reflected by different subpopulations with distinct gene expression. To preserve the developmental progression of these subsets of endothelial cells, we used PHATE (Potential of Heat-diffusion for Affinity-based Trajectory Embedding) to analyze our scRNA-seq data (Moon et al., 2017).

PHATE analysis demonstrated that *kdrl*+ mCherry cells formed a main vascular tree composed of 17 different branches corresponding to embryonic specification trajectories that were identified based on the expression of specific markers (Figure 2A). Based on the higher enrichment of artery and vein defining markers, we identified branches 1 and 2, as the main venous and arterial cell populations, respectively (Figure 2B). Branch 6 displayed co-expression of arterial and venous markers with cardiac genes, suggesting that this population mainly contributes to the heart vasculature. Branch 7 was enriched for markers of lymphatic endothelium, while the remaining cell pools expressed hemogenic, hematopoietic stem cells or blood progenitor markers (Figure 2B). Interestingly, *kdrl*+ mCherry cells isolated from mixed akt^Δ/Δ^ embryos showed an identical number of branches as those isolated from WT, further supporting that the loss of Akt did not grossly affect the patterning of the embryonic cardiovascular system (Figure 2C and D).

**Figure. 2.**
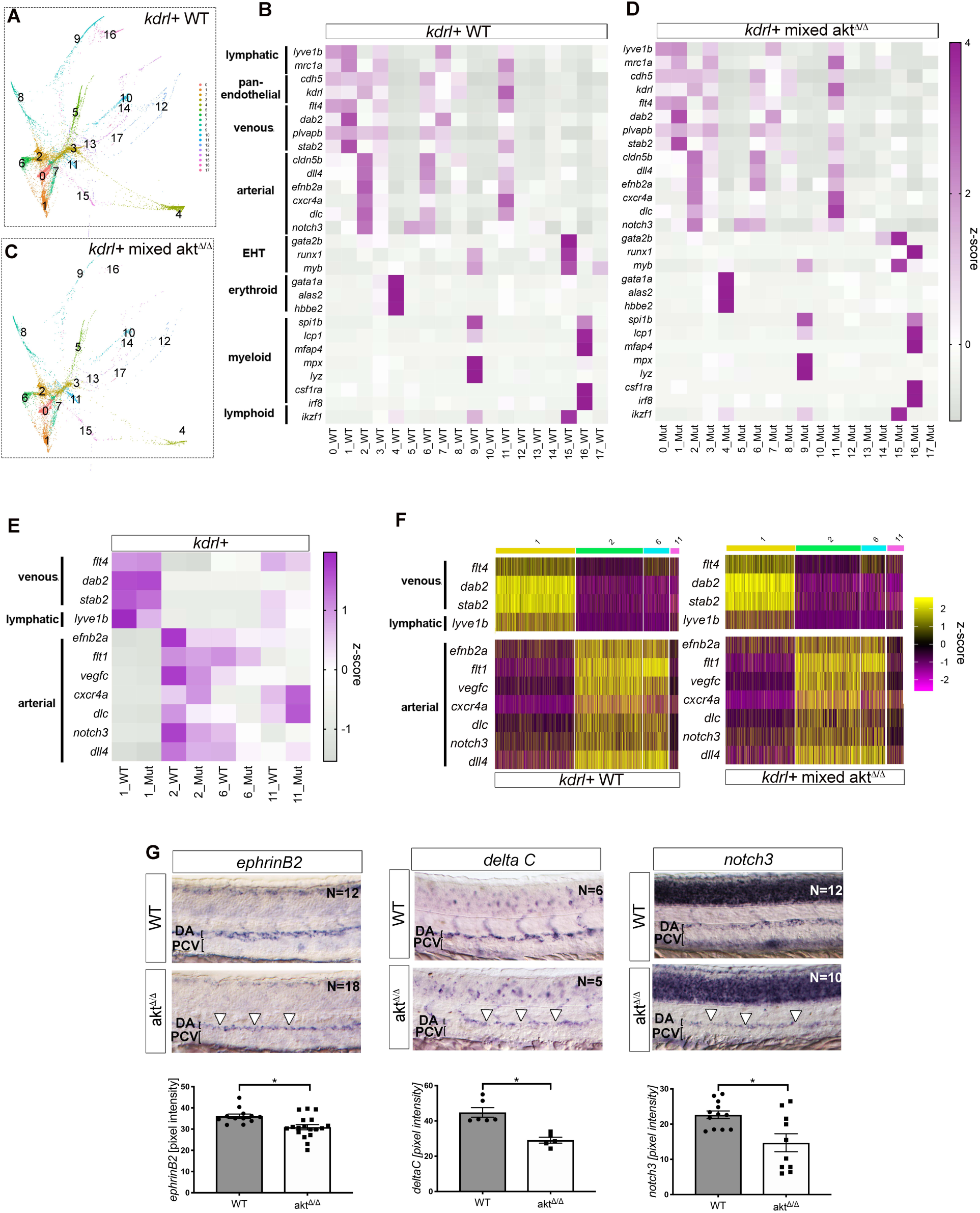
Single cell sequencing and in situ confirmation of diminished arterial markers expression in *akt*^*Δ/Δ*^ zebrafish. PHATE plots (A,C) and heat maps (B,D) of scRNASeq data from WT and mixed *akt*^*Δ/Δ*^ embryonic kdrl+ endothelial cells, respectively. All heatmaps depict z-score of vascular gene expression levels in kdrl+ endothelial cells expressing transcripts in scRNA-seq PHATE branches. (E) Heatmap highlighting the expression of venous, lymphatic and arterial markers from branches 1, 2, 6 and 11. (F) Heatmap showing increased fraction of cells in branch #11 in mixed *akt*^*Δ/Δ*^ embryos with the length of the line on the top delineating the fraction of cells within the endothelial cell population. (G) Whole mount *n situ* hybridization bright field images of arterial markers *ephrinB2, delta C* (*dlc*) and *notch3* in trunk vasculature of WT and *akt*^*Δ/Δ*^ embryos at 24 hpf. The embryos were placed in the lateral view with dorsal side up and anterior to left. Bar plots quantify pixel intensity of each marker. Data are mean ± SEM; *p< 0.05. DA=Dorsal Aorta, PCV=Posterior Cardinal Vein.

Next, we examined the differential expression of known vascular genes in the *kdrl*+ mCherry cell subsets from WT and mutant embryos. Strikingly, we observed diminished expression in mixed akt^Δ/Δ^ *kdrl*+ mCherry cells of genes that are mainly associated with artery differentiation and Notch regulated genes *(efnb2a, cxcr4, dlc, dll4, notch3*), but not venous fate (Figure 2E). Moreover, the fraction of cells within the branch 1 (vein) and 2 (artery) was reduced in the mutant endothelial cell population compared to WT, whereas the fraction of cells within the branch 11, which express both arterial and venous markers, was increased in the population of mutant endothelial cells as the expression of the arterial markers, *cxcr4* and *dlc* (Figure 2E and 2F). These data suggest that, at the molecular level, an excessive number of endothelial cells lacking *akt* fail to fully segregate into the arterial or/and the venous cell pool.

To verify the scRNA-seq results and add spatial resolution, we performed whole mount *in situ* hybridization of 24 hpf embryos to test the expression of artery markers *efnb2a, dlc* and *notch3* genes in the DA. Relative to WT embryos, *akt*^*Δ/Δ*^ embryos showed a decreased expression of all these arterial genes at 24 hpf (Figure 2G). In contrast, the expression of the venous marker, *ftl4*, or other markers for the maturation of spinal cord neurons (*ngn1, elav13*) or somites (*myoD*), showed no obvious changes in *akt*^*Δ/Δ*^ embryos versus WT (Figure S2). Thus, gene expression defects associated with altered arterial specification were specific and not the result of global developmental anomalies. Collectively, these data suggest that Akt signaling is required to sustain arterial endothelial cells specification.

### Akt drives artery specification independently of progenitor cell migration and ERK activity

To further investigate the mechanism(s) behind the vascular morphogenesis defects in *akt*^*Δ/Δ*^ embryos, we performed time lapse imaging from 20 to 22hpf in WT and mutant embryos. In WT embryos, endothelial progenitor cells sprouted from the DA primordium into the PCV at 20 hpf and fully segregated by 22-24 hpf (Figure 3A, left panel), as previously reported (Herbert et al., 2009b). Interestingly, in *akt*^*Δ/Δ*^ embryos, DA progenitors sprouted in greater numbers toward the PCV yet failed to completely separate within 22 hpf (Figure 3A, right panel). These data in *akt*^*Δ/Δ*^ embryos are reminiscent of loss of function *notch* mutants which manifest reduced Notch dependent gene expression (e*phrinB2, dl4, dlc)* in the DA (Lawson et al., 2002b; Quillien et al., 2014)(Figure 2F) or efnb2 morphants which result in the loss of artery specification and abnormal arteriovenous structures (Herbert et al., 2009a; Lawson et al., 2002b).

**Figure. 3.**
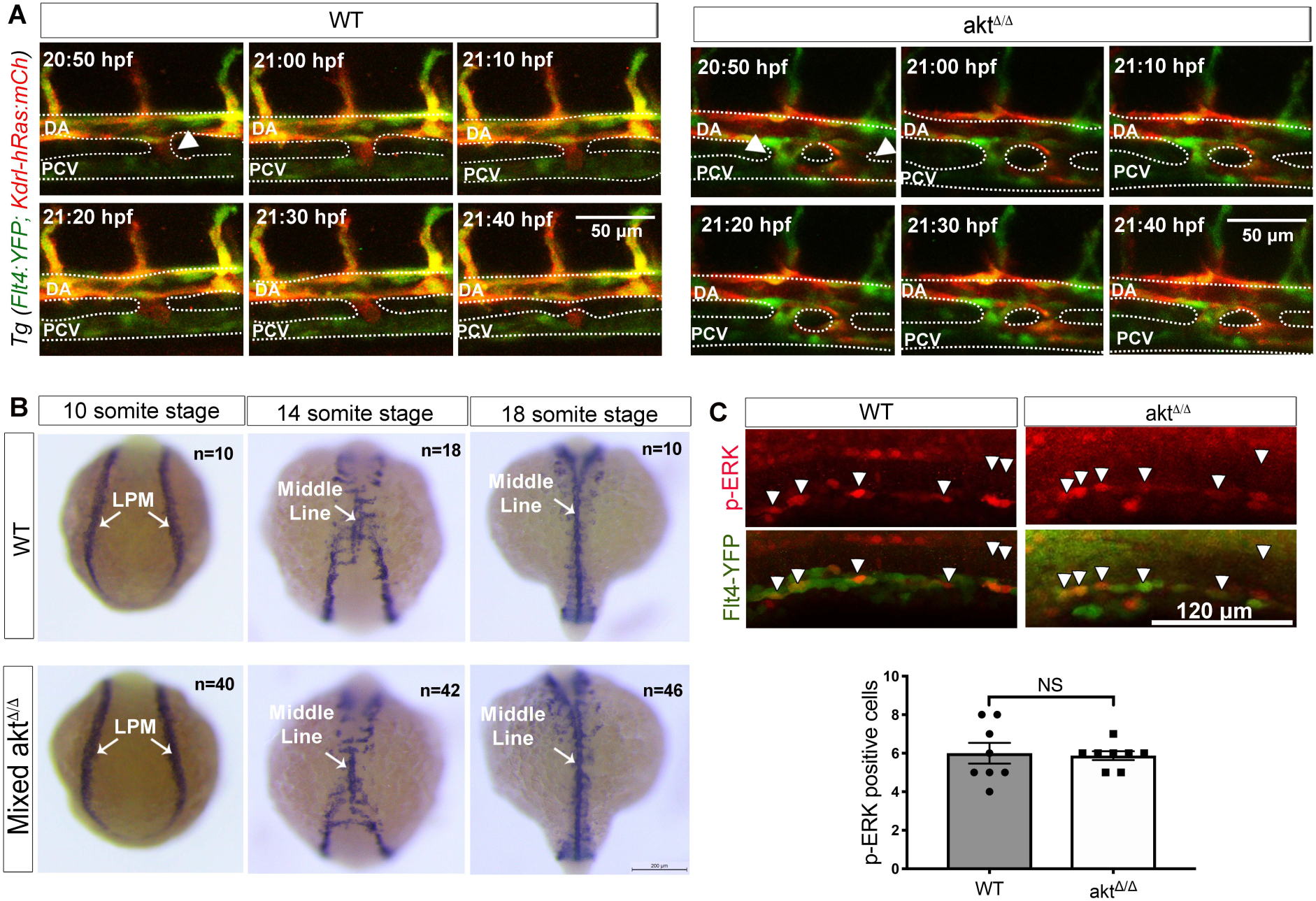
Loss of *akt* impedes arteriovenous separation, but not angioblast homing or ERK signaling. (A) Images from time lapse movies of WT and *akt*^*Δ/Δ*^ in *Tg (flt4:YFP; kdrl:hRas-mCherry)*^*hu4881;s896*^ embryos (20-22 hpf). The embryos were placed in the lateral view with dorsal side up and anterior to left. Timestamps are in upper left of each panel. White arrows indicate connection between the artery and vein. (B) *In situ* hybridization of marker *fli1a* labels the position of angioblasts at 10 ss, 14 ss and 18ss in WT and *akt*^*Δ/Δ*^ embryos. White arrows point to the lateral posterior mesoderm (LPM) at 10 ss and indicates the middle line formation at 14 ss and 18 ss. (C) Lateral view of immunofluorescence images showing phosphorylated ERK in DA of WT and *akt*^*Δ/Δ*^ in *Tg (flt4:YFP;)*^*hu4881*^ embryos at 14 ss. Data are mean±SEM. NS=non-significant.

The migration of endothelial progenitor cells from the lateral posterior mesoderm (LPM) toward the embryonic midline ensures vasculogenesis and artery specification via VEGFa-mediated regulation of ephrinB2 and Notch-signaling (Lawson et al., 2002a; Zhong et al., 2001). Given that Akt can be downstream of VEGFa, we investigated whether endothelial progenitor cells (*fli1a*+) in akt^Δ/Δ^ embryos have impaired migration from the LPM (Lawson and Weinstein, 2002b). *Fli1a*+ cells were visualized at different somite stages (ss) and no distinguishable differences were found between WT and mixed *akt*^*Δ/Δ*^ embryos from 10 to 18 ss (Figure 3B). Interestingly, the levels of somitic *Vegfa* expression at 18 ss were enhanced (Figure S3A) in *akt*^*Δ/Δ*^ versus WT embryos. Thus, the artery specification or altered arteriovenous separation defects in *akt*^*Δ/Δ*^ embryos were not a consequence of impaired Fli1a+ cell migration or reduced *Vegfa* expression. We also examined whether anomalies in cell survival could explain *akt*^*Δ/Δ*^ vascular phenotypes, but we did not observe significant differences in the number of apoptotic cells between WT and *akt* mutants (Figure S3B).

Previous studies suggested that ERK activation is required downstream of VEGFa for arterial identity and proliferation and that input from the PI3 kinase/Akt pathway may negatively regulate the extent of ERK activation (Hong et al., 2006; Ren et al., 2010). Notably, p-ERK was localized in endothelial progenitor cells of the DA primordium (Shin et al., 2016) however, the number of p-ERK+ cells was comparable in WT and *akt*^*Δ/Δ*^ 20 ss embryos (Figure 3C). Thus, our data suggest that Akt is distinctly required for specification of the main embryonic artery, independent of ERK-signaling.

### Akt control DA formation upstream FOXO and Notch signaling pathways

We found that the DA specification is severely affected in the Akt loss of function embryos, while the PCV specified normally. Interestingly, both vessels showed a reduction in vascular diameter likely due to the excessive number of cells failing to segregate either in the DA or PCV. To interrogate the signaling pathways upstream and downstream of Akt we therefore use DA morphology and arteriovenous (A-V) connections at 24hpf as the main phenotypic read-out of Akt loss of function. We first employed pharmacological agents that selectively interfere with components of the VEGF-PI3K pathway, which lie upstream in the Akt signaling cascade. Treatment of embryos with SU5416, a selective inhibitor the VEGF receptor 2 kinase (kdrl) (Kasper et al., 2017), decreased DA diameter in both WT and mixed *akt*^*Δ/Δ*^ embryos (Figure 4A, top for images and 4B for quantification). Interestingly, SU5416 enhanced A-V connections in WT but not further in mutant embryos indicating that VEGF signaling contributes to DA assembly and A-V separation but the effect on A-V connections is impeded in mixed *akt*^*Δ/Δ*^ mutants. Treatment with the PI3K inhibitor LY-294,002 (Chen et al., 2017) reduced DA diameter in WT embryos but did not further impact DA in *akt*^*Δ/Δ*^ embryos (Figure 4C, top for images and 4B for quantification). Similarly to SU5416 treated embryos, blockade PI3K increased A-V connections in WT but not further in mutant zebrafish (Figure 4B, top for images and below for quantification). These results indicate that VEGF-kdrl-PI3K signaling is critical for DA development and A-V separation, in part, via Akt.

**Figure. 4.**
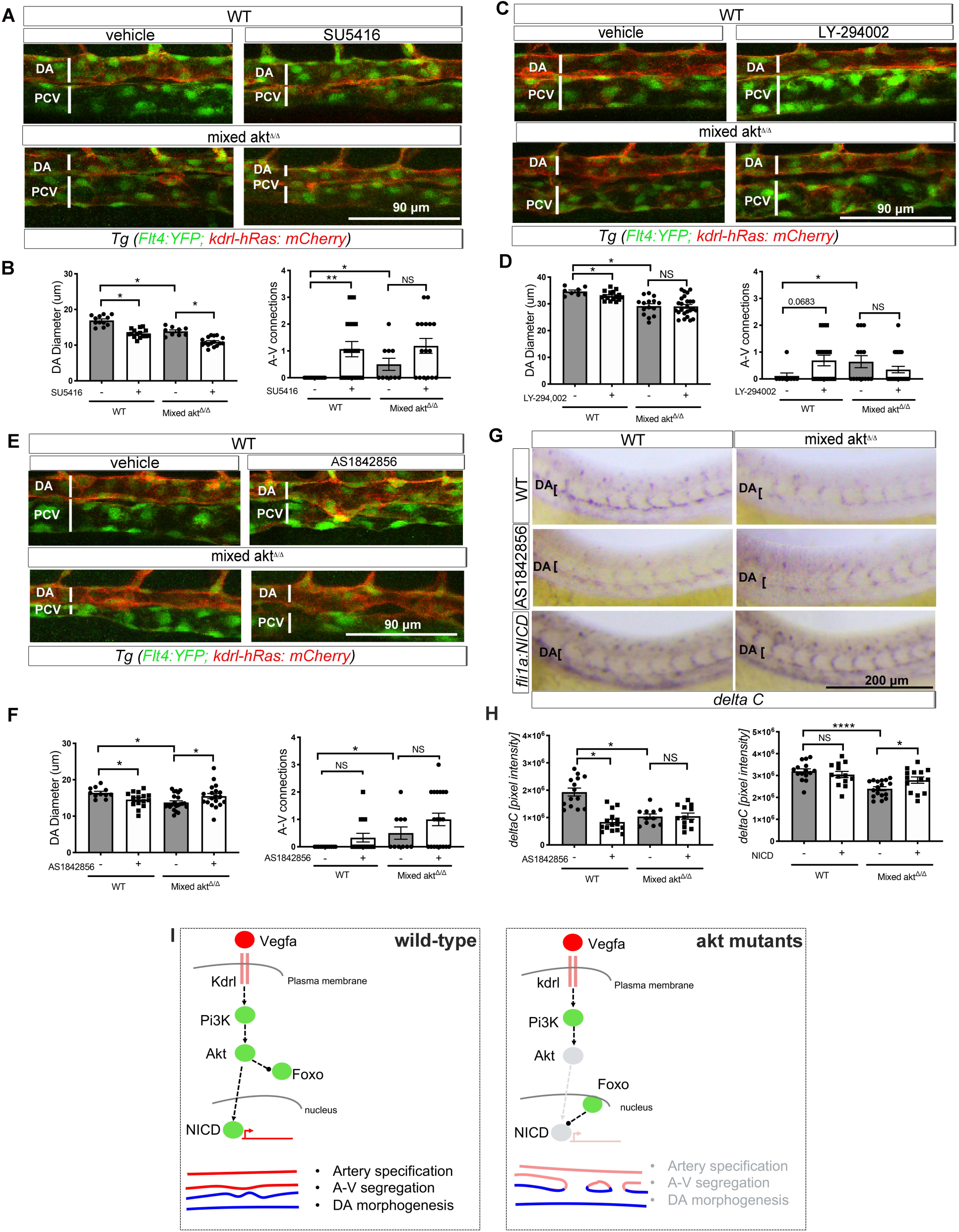
Loss of *akt* reduces DA diameter and arterial specification via FOXO and Notch. (A) Lateral view images and (B) quantification of DA diameters and A-V connections in the trunk vasculature after treatment of WT and mixed *akt*^*Δ/Δ*^ embryos at 24 hpf with a selective inhibitor of the VEGF receptor 2 (SU-5416, 1 μM). (C) Lateral view images and (D) quantification of DA diameters and A-V connections of the trunk vasculature after treatment of WT and mixed *akt*^*Δ/Δ*^ embryos at 24 hpf with a PI-3K inhibitor (LY-294,002, 80 μM) (E) Lateral view images and (F) quantification of DA diameters and A-V connections of the trunk vasculature after treatment of WT and mixed *akt*^*Δ/Δ*^ embryos at 24 hpf with a FOXO inhibitor (AS1842856, 0.1 μM). (G) *In situ* hybridization and (H) quantification of arterial marker *dlc* in WT and mixed *akt*^*Δ/Δ*^ embryos with and without treatment with FOXO1 inhibitor (AS1842856, 0.025 μM) and injection of an endothelia specific plasmid, *Fli1a:Notch-NICD*. Data are mean±SEM, * p<0.05, **p<0.01, ****p<0.0001. NS=non-significant. (I) Model for Akt signaling in WT and *akt* mutant zebrafish.

To investigate pathways downstream of Akt that may contribute to the vascular defects in mutant embryos, we investigated the transcription factor FOXO1, an established Akt substrate known to be critical for several aspects of angiogenesis (Dharaneeswaran et al., 2014). Akt phosphorylation of FOXO1 leads to its nuclear exclusion and alterations FOXO1 dependent gene expression. Notably, treatment with AS1842856, which binds and inhibits the non-phosphorylated, active form of the transcription factor FOXO1 in the nucleus (Gays et al., 2017) reduced the DA diameter in the WT embryos but increased DA diameter in mixed *akt*^*Δ/Δ*^ mutants (Figure 4E, top for images and 4F below for quantification), thus implying FOXO hyperactivation contributes to reduced DA in *akt*^*Δ/Δ*^ embryos (Wilhelm et al., 2016). However, FOXO did not contribute to A-V separation in WT or A-V in mutant embryos since inhibition of FOXO did not impact the number of A-V connections in any genotype, suggesting that FOXO contribute differently to DA formation and specification.

Since the loss of Akt reduces the levels of several Notch-dependent genes, *dll4, dlc*, and *notch3* in DA (Figure 2E), we assessed if FOXO can regulate the expression of the well characterized Notch dependent arterial specification marker *dlc*. Strikingly, inhibition of FOXO reduced the expression of *dlc* in the DA of WT embryos but did not further reduce the levels detected in *akt*^*Δ/Δ*^ embryos (Figure 4G, compare top and middle panels and 4H for quantification). These data suggest Akt phosphorylation of FOXO regulates *dlc* levels, arterial specification and DA morphology. However, in the Akt depleted state, FOXO blockage induces uncoupling of arterial specification (Figure 4H) from DA morphology (Figure 4F where FOXO inhibition rescues DA phenotype in mixed *akt*^*Δ/Δ*^ embryos).

Previous work has shown that Notch is downstream of Akt signaling (Bedogni et al., 2008; Kerr et al., 2016; Konantz et al., 2016; Zhang et al., 2011a). To verify if Notch is downstream of Akt during the DA specification, we expressed the Notch 1 intracellular domain (NICD) under the endothelial fli1a promoter to ectopically activate Notch signaling in WT and mixed *akt*^*Δ/Δ*^ embryos. Notably, NICD did not impact *dlc* expression in WT embryos, however NICD increased *dlc* levels in mixed *akt*^*Δ/Δ*^ embryos (Figure 4G, bottom panels and 4H for quantification in right graph). Thus, expression of NICD normalizes *dlc* expression in mixed *akt*^*Δ/Δ*^ embryos. These data suggest that Akt signaling to FOXO and Notch is critical for early arterial development and arterial specification.

## DISCUSSION

Akt is considered as an important signaling node co-opted by several angiogenic factors (VEGF, angiopoietin, ephrin B2, fibroblast growth factor) that integrates signal transduction mechanisms during angiogenesis and vascular homeostasis (Manning and Cantley, 2007). Despite this central role, the importance of Akt signaling during early vascular development has not been thoroughly addressed. Here, we show that the loss of *akt* in zebrafish impairs the formation of the first major vessel, the DA, by reducing arterial specification and arterio-venous segregation. Our single cell sequencing analysis of *akt* mutant endothelial cells support this phenotype. We found that the loss of Akt alters the size of distinct endothelial cell subpopulations, particularly those expressing arterial markers, and reduces the expression of several Notch-dependent genes as confirmed by *in situ* analysis. Similarly, Akt loss of function cells have an increased cell pool with arterial and venous co-expressed markers, thus fail to segregate within the DA and PCV vessels. Mechanistic analysis suggests that the loss of Akt did not impact the migration of endothelial progenitors towards the midline, the Vegfa availability or ERK activation but regulated the expression arterial fate markers dependent on FOXO and Notch (Figure 4I). Thus, these data place Akt at a critical signaling node during early artery-vein fate decisions in zebrafish.

Moreover, in Akt mutant embryos inhibition of FOXO increases the DA diameter to WT levels, while in presence of Akt, inhibition of FOXO has the opposite effect on the DA morphogenesis. This would suggest that FOXO function to control both DA diameter and specification is dependent on a threshold; namely that too high or too low FOXO activity results in similar DA defects. Another hypothesis is that FOXO controls DA morphogenesis and/or specification via parallel but independent mechanisms that might be antagonistic when the Akt-signaling pathway is lost (Figure 4I). Further studies will be required to characterize how FOXO function in Akt dependent and independent fashion.

Additionally, we show that Akt is required for expression of arterial endothelial cell specification genes independent of ERK, but dependent on Notch activity. Indeed, several Notch-dependent arterial markers were reduced in *akt*^*Δ/Δ*^ zebrafish, and overexpression NICD restored the expression of *dlc*, suggesting that Akt impacts Notch dependent arterial specification. Akt can regulate Notch activity via a direct or indirect function. For example, Akt functions upstream of Notch in aortic endothelial cells (Konantz et al., 2016), can directly phosphorylate Notch4-ICD (Lee et al., 2014) and regulate Notch4 nuclear localization (Ramakrishnan et al., 2015). Alternatively, angiopoietin-1 can activate Akt and promotes Notch signaling indirectly by phosphorylating glycogen synthase kinase 3β (GSK3β), thereby enhancing β-catenin activity and upregulating Dll4-Notch in endothelial cells (Lawson and Weinstein, 2002a; Zhang et al., 2011b). Furthermore, FOXO and Notch can directly interact to coordinate cell specification and function in the context of other tissues (Kitamura et al., 2007; Pajvani et al., 2011). Thus, it is possible that arterial cell specification is mediated by direct regulation of the Akt-FOXO-Notch pathway (Figure 4I).

Interestingly, single cell RNA sequencing of endothelial cells from embryos lacking Akt reveals an increase fraction of cells that co-express arteriovenous genes that do not segregate within classic artery and vein trajectories. The expression of genes controlling cellular responses to stress (e.g.: the myeloperoxidase *mpx* and a regulator of ER-mitochondrial contacts, Nogo-B) were decreased in this cell pool from *akt*^Δ*/*Δ^ embryos compared to WT(Astern et al., 2007) (Acevedo et al., 2004; Sutendra et al., 2011; Yu et al., 2009). Further studies will be necessary to investigate if and how genes within the arteriovenous unsegregated cells are directly regulated by Akt or if this defect is the result of Akt-Notch and/or Akt-FOXO-Notch regulation.

In summary, the genetic loss of Akt in zebrafish impairs arterial endothelial cell specification providing novel insights into the important role of Akt signaling pathway during early vascular development.

## Acknowledgements

This work was supported by Grants R01HL130246 (to SN), R35 HL139945, the Leducq Foundation (MIRVAD network), P01 HL1070205, AHA MERIT Award (to WCS), AHA Predoctoral fellowship (to WZ) and by the Swedish Cancer Foundation, Science Council and Knut and Alice Wallenberg Foundation (to CB).

We thank Meredith Cavanaugh for fish husbandry, Amelia Luciano, Ana Gamez, Andew Kuo, Ban Edani, Bo Tao, Dionna M. Kasper, EonJoo Park, Iwakiri Yasuko, Jan Kraehling, Jared Hintzen., Joseph Fowler, Kariona Grabinska, Liana Boraas, Lily Cheng, Nabil Boutagy, Monica Lee and Sungwoon Lee for discussing about the project, Madahiro Shin, Nathan Lawson, and Arndt Siekmann for providing fish line, Guilin Wang from Yale Center for Genome Analysis for scRNA seq data analysis.

## Author Contributions

WZ, SN, WCS designed experiments. WZ, ER, RZ and JJG performed experiments and/or analyzed the data. LH, SM and CB assisted with single cell sequencing analysis. WZ, SN and WCS wrote the manuscript. All authors contributed to the review of the manuscript.

## STAR METHODS

### KEY RESOURCES TABLE

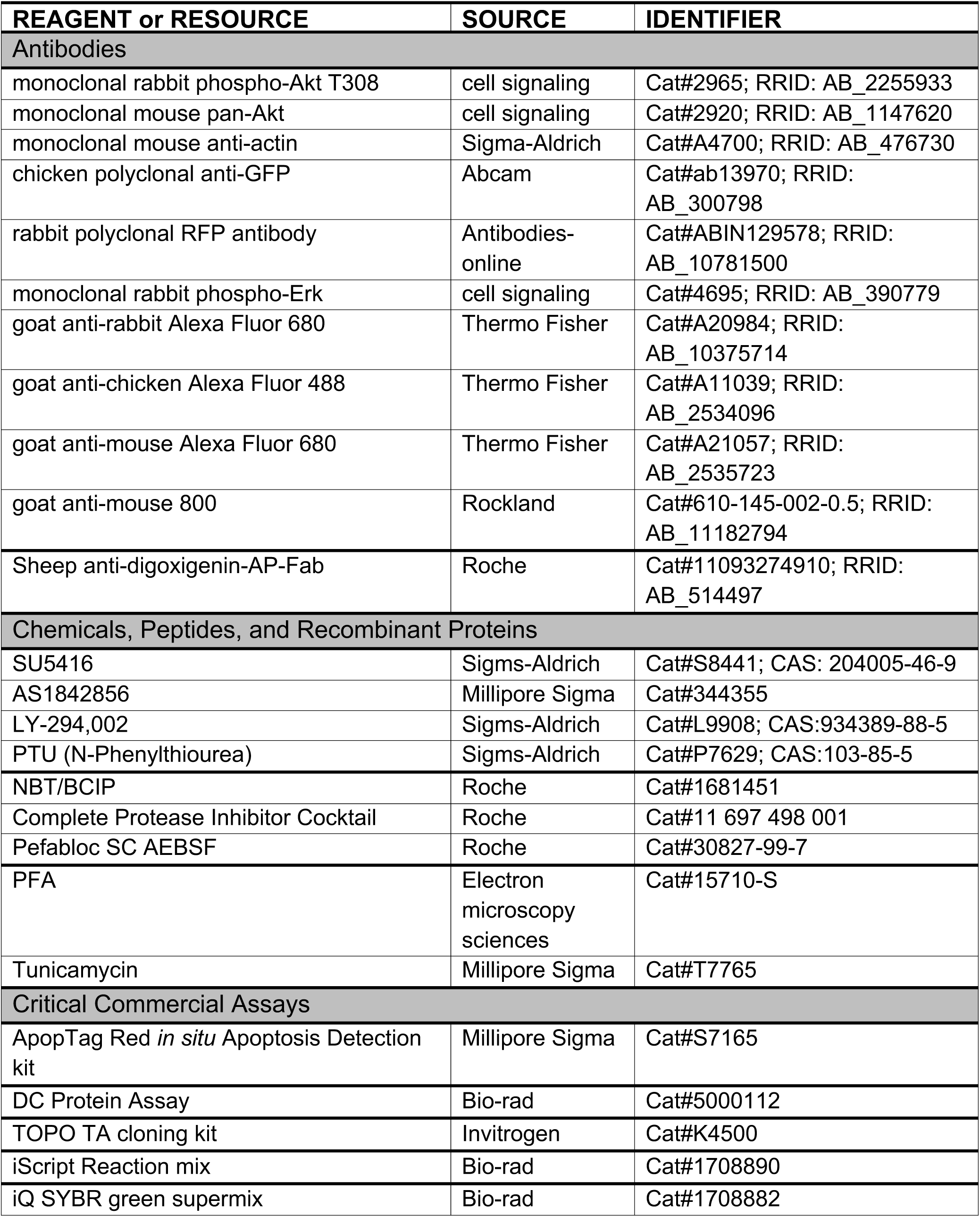
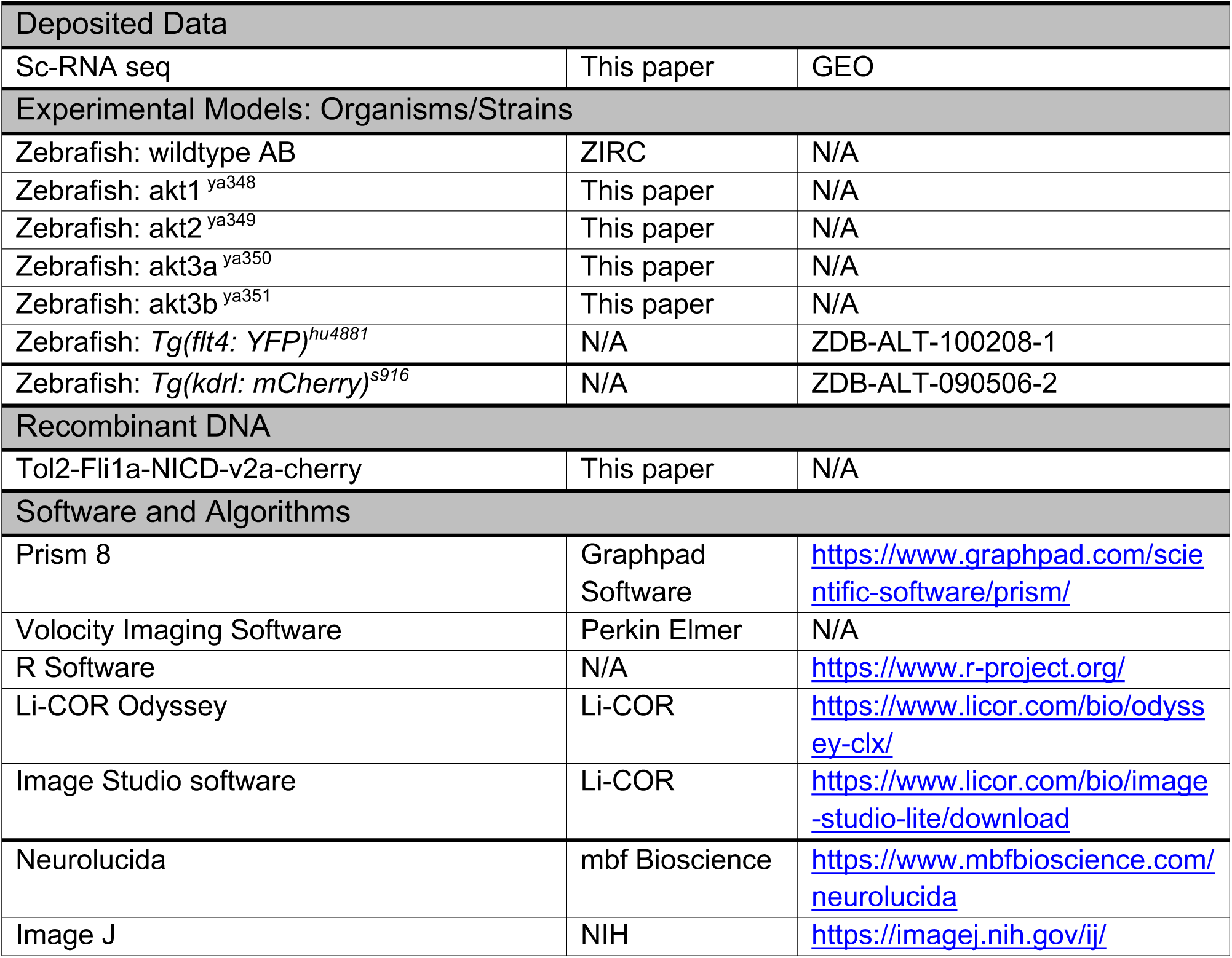

### LEAD CONTACT AND MATERIALS AVAILABILITY

Requests for information and materials can be sent to Stefania Nicoli and William Sessa.

### EXPERIMENTAL MODEL AND SUBJECT DETAILS

#### Generation of Akt loss-of-function zebrafish and zebrafish husbandry

Zebrafish were raised and maintained at 28.5°C using standard methods (unless otherwise indicated), and according to protocols approved by Yale University Institutional Animal Care and Use Committee (# 2017-11473).

gRNA generation & injection: CRISPRScan (Moreno-Mateos et al., 2015; Vejnar et al., 2016) was used to design gRNAs to mutate the Akt 1, 2, 3a 3b loci with CRISPR/Cas9 genome editing (sequence Table S1). The gRNA template for in vitro transcription was PCR amplified and purified with Qiaquick PCR purification Kit (Qiagen). In vitro transcription followed and was performed as described (Narayanan et al., 2016).. F0 *Tg(flt:YFP; kdrl: mCherry)*^*hu4881s916*^embryos injected with 100 pg of gRNA and 200 pg of Cas9 mRNA at the one-cell stage were used to confirm mutagenesis via T7 endonuclease I (T7E1) assay and sequencing as described (Narayanan et al., 2016) or were analyzed in phenotypic assays. Genomic DNA was isolated from a clutch of injected embryos with 100 mM sodium hydroxide and 1M pH 7.5 Tris-HCl and then amplified regions including the mutation site using PCR. After 3 months, the F0 founder fish was genotyped using fragment analysis (DNA analysis Facility Yale University) and out-crossed with WT (sequence Table S1). F1 allele were identified, genome PCR product were cloned into TOPO vector with the TOPO TA cloning kit (Invitrogen) for sequencing. The sequencing results showed that the mutant alleles carried the following out loss of function mutation: Akt1 ^ya348^ (−1 based pair deletion), Akt2 ^ya349^ (+10 based pair insertion), Akt3a ^ya350^ (−21 based pair deletion) and Akt3b ^ya351^ (−4 based pair deletion), which are labeled as homozygous *akt1*^*Δ/Δ*^, *akt2*^*Δ/Δ*^, *akt3a*^*Δ/Δ*^ and *akt3b*^*Δ/Δ*^ in the paper. The mixed *akt*^*Δ/Δ*^ in this study were generated from the in-cross of akt1, akt3a, akt3b homozygous and akt2 heterozygous mutant.

For DNA injection, embryos were injected in the one-cell with 25 pg of the expression construct Tol2-Fli1a-NICD-v2a-cherry and Tol2 transposase mRNA, and later selected for mCherry expression.

### METHOD DETAILS

#### Single cell RNA sequencing

WT and Akt loss-of function zebrafish embryos were collected at 24 hpf, dechorionated and placed in 2 ml tubes with egg water. Embryos were disassociated into single cell suspensions and subjected to Fluorescence Activated Cell Sorting (FACS) as described for transgenic zebrafish previously (Ristori and Nicoli, 2015), *Tg(flt4:YFP; kdrl:mCherry)*^*hu4881s916*^ endothelial cells were FAC-sorted into 0.04% BSA in 1x PBS and the captured endothelial cell fraction was loaded onto the 10X Genomics Chromium instrument for a targeted recovery of 10,000 cells per sample. The reaction was then reverse transcribed and barcoded using the 10X Genomics Chromium Next GEM Single Cell 3’ Library Construction Kit v2.1 according to manufacturer instructions.

The barcoded libraries were sequenced on Illumina HiSeq 4000 instrument. The reads were de-multiplexed using 10X Genomics provided version of program bcl2fastq. These fastq files were further processed with cellranger and indexed custom genome for *D. rerio* with sequences for mCherry. The output of cellranger program was processed using the package Seurat (Satija et al., 2015; Stuart et al., 2019) in R (Team, 2019). Briefly, the data was filtered such that all the cells that had less than unique transcripts detected or had more than 10% of the transcripts coming from mitochondrial genes were eliminated, as were the genes that were seen in less than 3 cells. Filtered data was then normalized and scaled using sctransform (Hafemeister and Satija, 2019). The data was further clustered in Seurat, and then rendered as a PHATE plot (Moon et al., 2019).

#### Quantitative RT-PCR

Embryos were processed whole or disassociated into single cell suspensions and subjected to FACS as described above for zebrafish scRNA-seq. 300 ng RNA was mixed with iScript Reaction mix and reverse Transcriptase (Bio-Rad) to generate cDNA with a 20 ul reaction (Kasper et al., 2017). qRT-PCR primers are listed in Table S1.

The 2^−ΔCT^ or 2^−ΔΔCT^ method was used to determine relative gene expression for quantitative RT-PCR analyses as indicated. mRNA levels were normalized to the beta actin housekeeping gene, *actb1* and was relative to the indicated control. Statistical comparisons between replicate pair ΔCT values for indicated groups were determined by a paired, two-tailed Student’s t-test.

#### Western Blot

Protein extraction was performed as following: 10-20 WT and Akt loss-of function zebrafish embryos (4 days post fertilization) were placed in 1.5 ml tubes and 1x PBS. Embryos were pipetted with 20 ul tip to break the zebrafish tissues and centrifuged at 1300 rpm for 10 minutes. After removing the supernatant, pellets were re-suspended in the lysis buffer with 2 ul per embryo. The lysates were left on ice for 15 minutes, mixed for 15 seconds and span at 12000 rpm for 10 minutes at 4 C. The supernatant was transferred to new cold tubes, sonicated for 6 seconds and protein concentration measured using DC Protein Assay (Bio-rad). Samples were mixed with 6x protein buffer and boiled for 5 minutes. About 20 ug protein samples were added into 10% SDS– polyacrylamide gel electrophoresis (SDS-PAGE) well and run at 80 V for 20 minutes and 100 V for about 1 hour. Then the SDS-PAGE was transferred to 0.45-μm nitrocellulose membranes (Bio-Rad) at 100 V for 2 hours. The nitrocellulose membranes were by 5% Bovine serum albumin (BSA) blocking buffer for 1 hour, incubated with primary antibody at 4°C overnight, washed with TBST (Tris-buffered saline, 0.1% Tween 20) for 5 minutes 3 times, incubated with and washed with TBST for 5 minutes 3 times. Blots were visualized by Li-COR Odyssey and analyzed by the Image Studio software (Li-COR).

Lysis buffer included 50 mM Tris-HCl, 1% NP-40 (v/v), 0.1% SDS, 0.1% Deoxycholic acid, 0.1 mM EDTA, 0.1 mM EGTA and 8 mg NaF, 4 mg sodium pyrophosphate, 20 mg Complete Protease Inhibitors (Roche), 3 mg Pefabloc SC AEBSF (Roche), 0.25 mM Sodium orthovanadate and 50 mM β-glycerolphosphate in 10 ml lysis buffer. Protein buffer contains 70 ml Tris-HCl, 36 ml glycerol, 10 g SDS, 6 ml 2-Methylbutane, 40 mg bromphenol-blue and water to 120 ml. Primary antibodies (1:1000) were monoclonal rabbit phospho-Akt T308 (cell signaling #2965), monoclonal mouse pan-Akt (cell signaling #2920) and monoclonal mouse anti-actin (Sigma-Aldrich). Secondary antibodies (1:10,000) were goat anti-rabbit Alexa Fluor 680 (Invitrogen), goat anti-mouse Alexa Fluor 680 (Invitrogen) and goat anti-mouse 800 (Rockland).

#### Whole mount in situ hybridization

Whole mount in situ hybridization (WISH) was performed using hybridization probes previously described (Kasper et al., 2017). Images of at least 6-12 stained embryos per replicate were blinded and quantified as follows. Briefly, images were converted to 8-bit and processed with FFT bandpass filter, filtering large structures to 200 pixels. Region(s) of interest were identified via thresholding and staining area or cluster number was measured using Analyze Particles(Dobrzycki et al., 2018b). Area measurements or cluster counts from biological replicates per genotype were combined and an unpaired, two-tailed Mann-Whitney U test was applied to detect statistical differences between conditions.

#### Immunofluorescence

WT and Akt loss-of-function embryos (24 hpf) were fixed and washed with the same condition as performed in WISH. After permeabilization with 0.125% Trypsin in PBS on ice for 20 minutes, samples were washed with PBST for 5 minutes 3 times and blocked at 4°C for 3-4 hours. Primary antibodies were monoclonal rabbit phospho-Erk (cell signaling #4695) (1:100), chicken polyclonal anti-GFP (1:300) (Abcam) and rabbit polyclonal RFP antibody (antibodies online) (1:300) were mixed with samples overnight. Samples were washed 6 times 45 minutes each and incubated with secondary antibodies (invitrogen) (1:300) goat anti-rat Alexa Fluor 594, goat anti-chicken Alexa Fluor 488 and goat anti-rabbit Alexa Fluor 633 overnight and washed with PBSTw 6 times each 45 minutes. Embryos were imaged with Lecia SP8 microscope.

#### TUNEL assay

30-40 WT and Akt loss-of-function zebrafish were placed in 1.5 ml tubes and fixed with 4% PFA at 4°C overnight. Samples were washed with PBS for 3 times, placed in MeOH at -20°C overnight and washed with 75% MeOH/PBST, 50% MeOH/PBST, 25% MeOH/PBST and PBST for 5 minutes respectively. After treatment with 10 ug/ml proteinase K at room temperature for 1.5 minutes, samples were washed with PBST twice and re-fixed in 4% PFA at room temperature for 20 minutes. Samples were washed with PBST for 5 minutes 5 times and incubated with pre-chilled Ethanol:Acetic Acid (2:1) at -20°C for 10 minutes. Followed with 3 times wash of PBST for 5 minutes each, samples were incubated at room temperature in 75 ul equilibration buffer for 1 hour (ApopTag Red *in situ* Apoptosis Detection kit (Millipore)). A small volume of working strength TdT (70% reaction buffer and 30% TdT with 0.3% Triton 100) was added into the reaction and incubated at 37°C overnight. The positive control was incubated with DNAseI overnight. The negative control was without TdT. The reaction was stopped with working strength stop/wash buffer (1 ml concentrated buffer with 34 ml dH_2_O) for 3-4 hours at 37°C. After wash with PBST 3 times for 5 minutes, samples were blocked with 2 mg/ml BSA, 5% sheep serum in PBST for 1 hour at room temperature and incubated in dark for 45-60 minutes at room temperature with working strength rhodamine antibody solution. After 30 minutes wash with PBST for 4 times, samples were incubated for 15 minutes with DAPI (1:200) in PBST and washed 3 times with PBST.

#### Chemical treatments

10 hpf zebrafish embryos were treated with 0.025 μM FOXO1 inhibitor (AS1842856, Millipore Sigma), 1 μM Sugen 5416 (SU5416, Sigma-Aldrich), 80 μM PI3K inhibitor (LY-294,002, Sigma-Aldrich). At 24 hpf treated embryos were fixed an processed for immunofluorescence or whole mount in situ hybridization.

#### Image Acquisition

Zebrafish embryos imaged by confocal or bright-field microscopy were raised in 0.003% 1-phenyl-2-thiourea (PTU) starting after the gastrulation stage to prevent pigmentation. Embryos imaged live by confocal microscopy were anesthetized in 0.1% tricaine and mounted in 1% low melt agarose. The majority of fluorescent images and time lapse movies were captured with an upright Leica Microsystems SP8 confocal microscope. Confocal time lapse movies were performed at room temperature starting at ∼20 hpf with z-stacks acquired at an interval of 11 minutes for a total of 15 hours. Bright field images of whole mount in situ staining were acquired with a Leica Microsystems M165FC stereomicroscope equipped with Leica DFC295 camera.

### QUANTIFICATION AND STATISTICAL ANALYSIS

#### Live fish imaging quantification

Using the software Imaris, DA, PCV and ISV were tracked manually and information of DA and PCV diameters and ISV length quantified and used for statistical analysis.

#### Statistical analysis

The quantification of *in situ* hybridization is followed the method of the Monteiro group (Dobrzycki et al., 2018a). The difference is that I used the nonparametric t-test (Mann-Whitney test) instead of 2-tailed ANOVA test. The other quantification is also nonparametric t-test (Mann-Whitney test). Z-scores were calculated based on the average gene expression and standard deviation across all PHATE branches for an individual gene.

### DATA AND CODE AVAILABILITY

The scRNA sequencing results were submitted to Gene Expression Omnibus.

## SUPPLEMENTAL INFORMATION

There are four supplement figures, two movies and one table.

## SUPPLEMENTAL INFORMATION FOR

### Supplemental figures

**Figure. S1.**
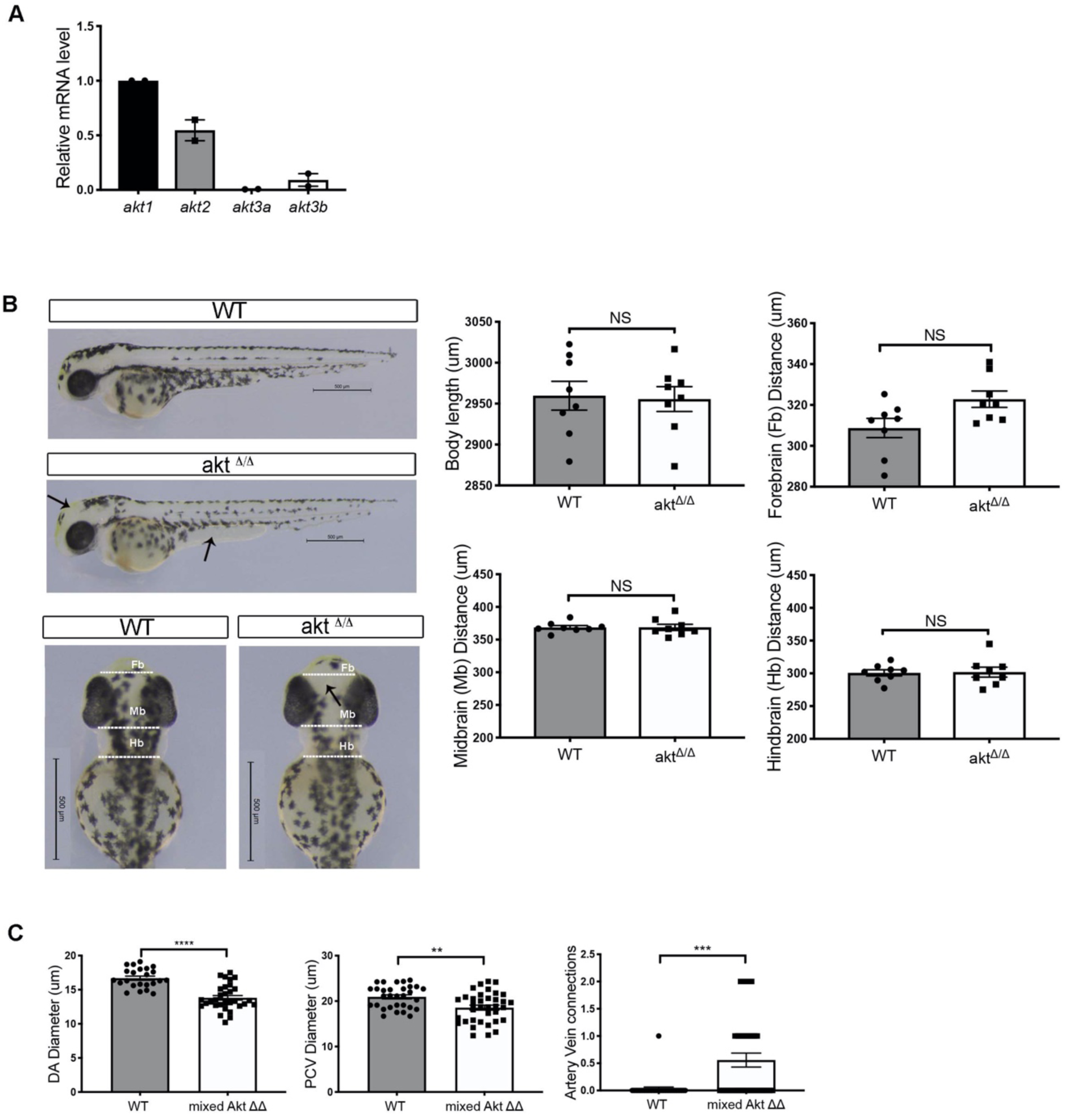
*Akt* ^*Δ/Δ*^ *embryos exhibit normal morphology*. (A) *akt1, 2, 3a* and *3b* mRNA levels in endothelial cells at 24 hpf. Endothelial cells were isolated from *Tg (kdrl:mcherry)* using FACS. (B) Bright field images of the body and head morphology of WT and *akt*^*Δ/Δ*^ embryos at 48 hpf and quantification on right side. Black arrows demonstrate the less pigment in the embryos; doted white line indicate the width of brain regions. (C) Bar plots show in WT, and *mixed akt*^*Δ/Δ*^ cardiovascular defects as reported in Figure 1C for *akt*^*Δ/Δ*^ embryos. Data are mean ± SEM for DA and PCV diameters and artery and vein connections; **p< 0.01, ***p<0.001, ****p<0.0001. DA=Dorsal Aorta, PCV=Posterior Cardinal Vein. Fb=Forebrain, Mb=midbrain, Hb=Hindbrain.

**Figure. S2.**
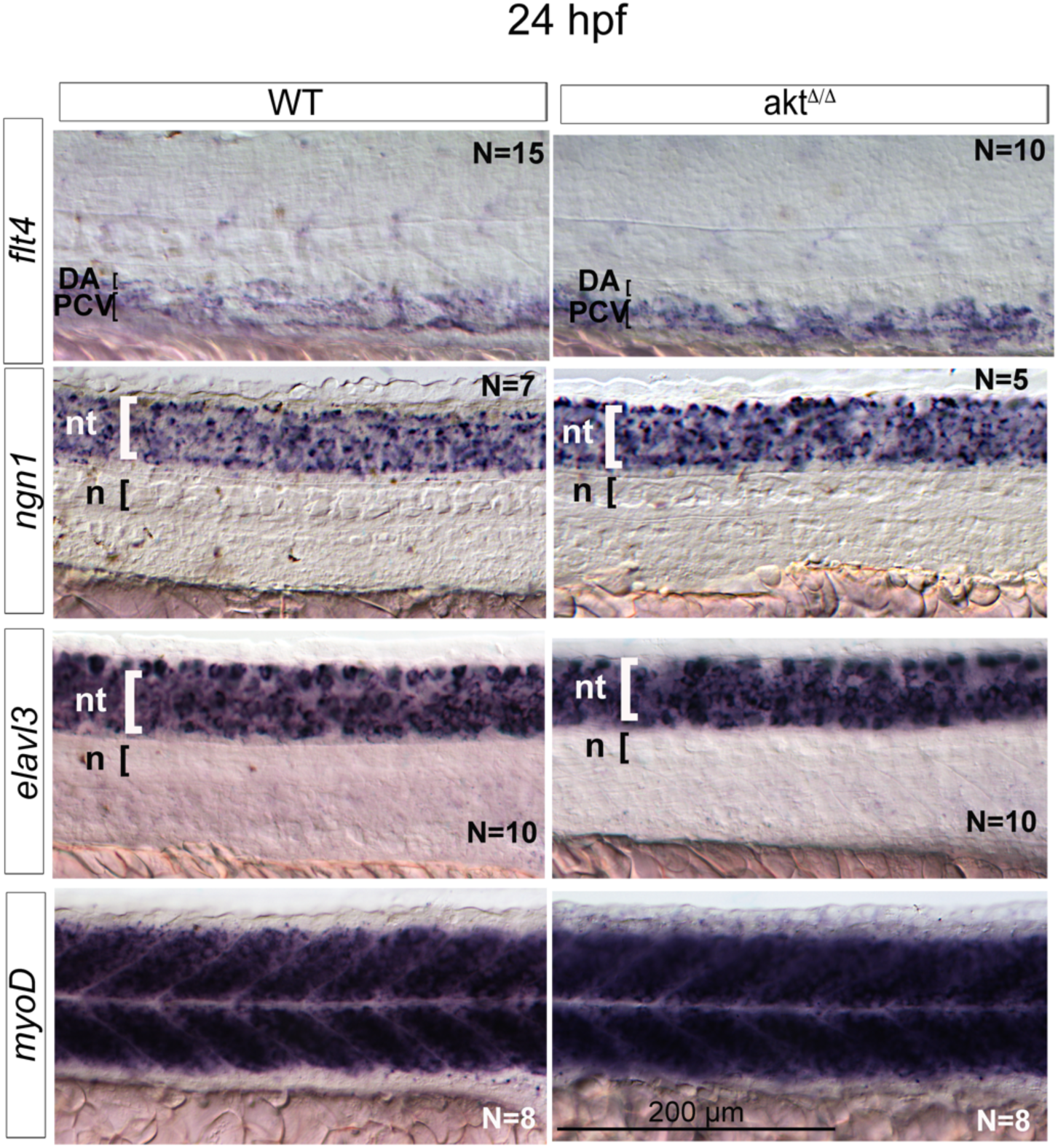
*akt*^*Δ/Δ*^ embryos express normal venous and non-vascular specification markers. *Whole mount situ* hybridization images show flt4 venous marker, mature neuron markers *ngn1* and *elavl3* and somite marker *myoD* in WT and *akt*^*Δ/Δ*^ embryos at 24 hpf trunk. All images represent the lateral view of zebrafish embryos, head to the left. nt: neural tube, n: notochord, DA=Dorsal Aorta, PCV=Posterior Cardinal Vein.

**Figure. S3.**
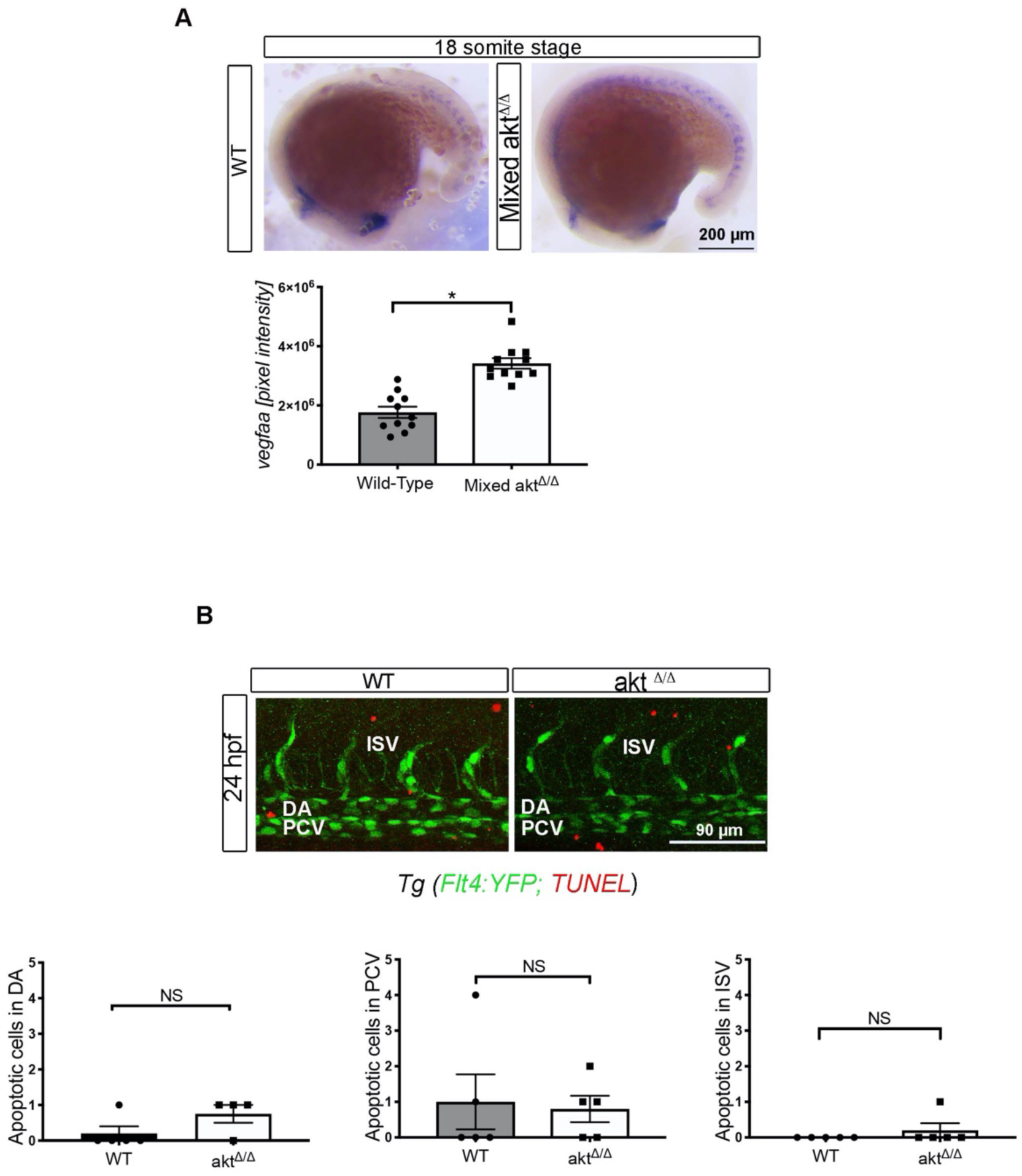
Akt drives DA specification without inhibiting vegfa, flow or promoting cell death. (A) Bright field images of whole mount *in situ* hybridization showing *Vefgaa* expression at 18ss and quantification. (B) Confocal image of WT or *akt*^*Δ/Δ*^ embryos at 24 hpf trunk with TUNEL assay labeling the apoptotic cells and quantification. All images represent the lateral view of zebrafish embryos, head to the left. Data are expressed as mean±SEM, *p<0.05 and NS=non-significant. DA=Dorsal Aorta, PCV=Posterior Cardinal Vein, ISV=Intersegmental vessels.

### Supplemental table

**Table S1.**
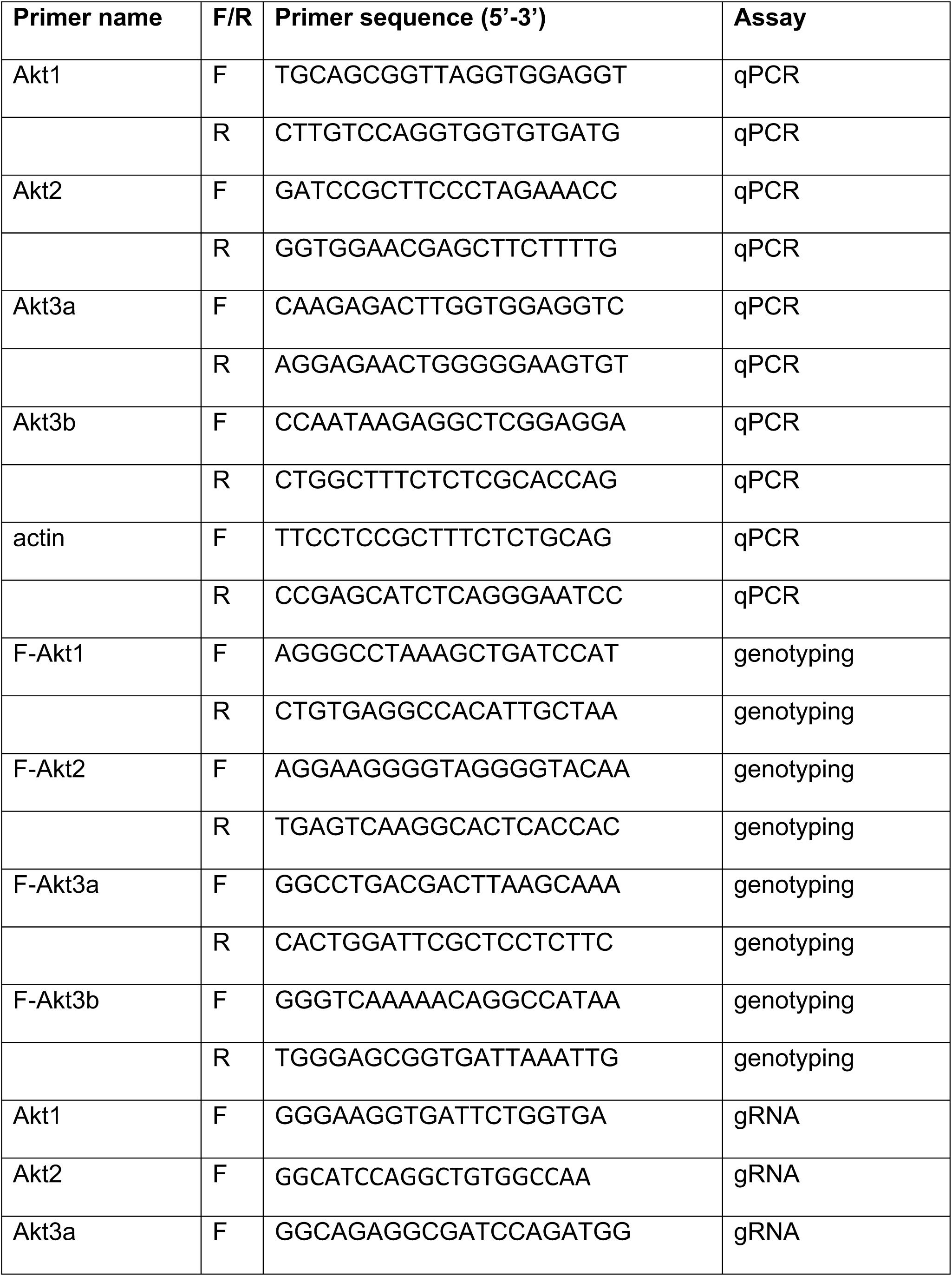
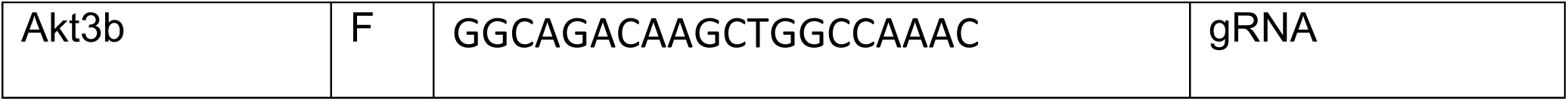
gRNA, qPCR and genotyping primers used in the paper.

